# Characterizing top-down microcircuitry of complex human behavior across different levels of the visual hierarchy

**DOI:** 10.1101/2022.12.03.518973

**Authors:** Logan Dowdle, Geoffrey Ghose, Steen Moeller, Kamil Ugurbil, Essa Yacoub, Luca Vizioli

## Abstract

FMRI has become a key tool for human neuroscience. At ultra-high field (=> 7 Tesla) it is possible to acquire images with submillimeter spatial precision, which allows examinations of mesoscale functional organization. Studying the brain at this scale does not come without its challenges, however. To tackle some of these challenges, we propose an approach that builds upon task modulations to identical visual stimuli and the simultaneous imaging of distal areas of varying complexity across the cortical hierarchy. Specifically, we record BOLD responses elicited by face stimuli during a stimulus relevant and a stimulus irrelevant task across cortical depths in V1, Occipital Face (OFA) and Fusiform Face area (FFA). We observed that task-related top-down modulations were larger in the inner compared to the outer layers of V1; and in the outer compared to the inner layers in the FFA. Our findings are consistent with animal reports of feedback exchange between deeper and superficial layers and with the notion of apical dendritic amplification as a key mechanism of conscious perception. Our approach showcases the potential of “laminar-fMRI” to explore large scale network activity and represents a promising step towards characterizing laminar functional profiles in humans for complex, cognitively meaningful, and socially relevant stimuli such as faces.

## Main

Since its inception 30 years ago^1,2^, functional MRI (fMRI) has become one of, if not the main tool for human neuroscience. This successful evolution is in part due to its non-invasiveness, relatively high spatial precision, and the potential for whole-brain coverage. Recent advances in MRI hardware and software have enabled the acquisition of functional images with unprecedented temporal and spatial resolutions in humans^3–7^. This allows, at least in principle, studying some fundamental units of neural computation - cortical layers and columns^5,8–10^. Until recently, access to such mesoscopic organizations had been accessible only by means of invasive electrophysiology in animals. Animal studies have shown that cortical networks are characterized by a specific laminar architecture, with feedforward/bottom-up signals mainly arriving in the middle layers, and feedback/top-down signals in the outer and inner layers^11–13^. Given the intricacy of the human brain, however, which has been parcellated into more than 300 regions compared to the ~50 cortical areas identified in non-human primates, animal models may fail to accurately describe the hierarchical complexity of human laminar processing. The need for human specific laminar studies is therefore self-evident.

The non-invasive nature of submillimeter fMRI, in conjunction with its potential for large cortical coverage and the ability to modulate the amount of top-down signal by varying task demands^14^, offers the unique possibility to characterize human mesoscale functional organization. Submillimeter acquisition protocols, however, operate in an SNR-starved thermal noise dominated regime; they inherently have low SNR due to smaller voxel volumes and incur in further losses in SNR from acceleration. As such, most human laminar studies are limited in coverage and confined to early sensory cortices that are less challenging to image (for example, V1, where signal and responses are large, or M1, where the cortex is thickest). However, in order to fully realize the potential and promises of laminar fMRI, researchers must move beyond early sensory cortices and also examine these areas in conjunction with higher-level regions^15^.

To address these challenges, we introduce both acquisition and analysis innovations. We record BOLD responses at 7T elicited by faces using an fMRI voxel volume of 0.343 μL (0.7mm isotropic resolution), which represents approximately a 33% decrease relative to the 0.512 μL typically employed in human mesoscale studies, while concomitantly increasing cortical coverage, which allowed the simultaneous imaging of visual and ventral cortices. We were able to achieve this milestone via the use of our recently developed NORDIC denoising^16,17^ technique which suppresses thermal noise without a meaningful impact on spatial resolution or related point spread function. Crucially, to examine top-down modulations on the BOLD signal, we implemented task manipulations, which we have recently shown to directly impact the responses elicited by identical visual stimuli^14^. This measure further allowed comparison of the signals acquired with identical stimulation but under different tasks constraints *within* cortical depths, and then compare these task effects across depths.

Specifically, participants viewed face images at varying levels of phase coherence (effectively manipulating the visibility of the stimuli), presented in a 12 seconds on/off block design fashion, while high resolution fMRI images of the occipital and ventral temporal cortices were collected (Figure 1. See Methods). For each task we collected 6 runs (with 8 blocks of stimuli per run, Figure 1) of high-resolution fMRI data. In each run, participants were instructed to perform one of two tasks – a *stimulus irrelevant* fixation task in which they responded to the fixation cross turning red; and a *stimulus relevant* face detection task in which they indicated whether they perceived a face. Importantly, to minimize interpolation induced blurring, fMRI data remained in their original, distorted space (see Methods) and upsampled to 0.35mm isotropic resolution, simultaneously with the motion correction step^18^.

**Figure 1.**
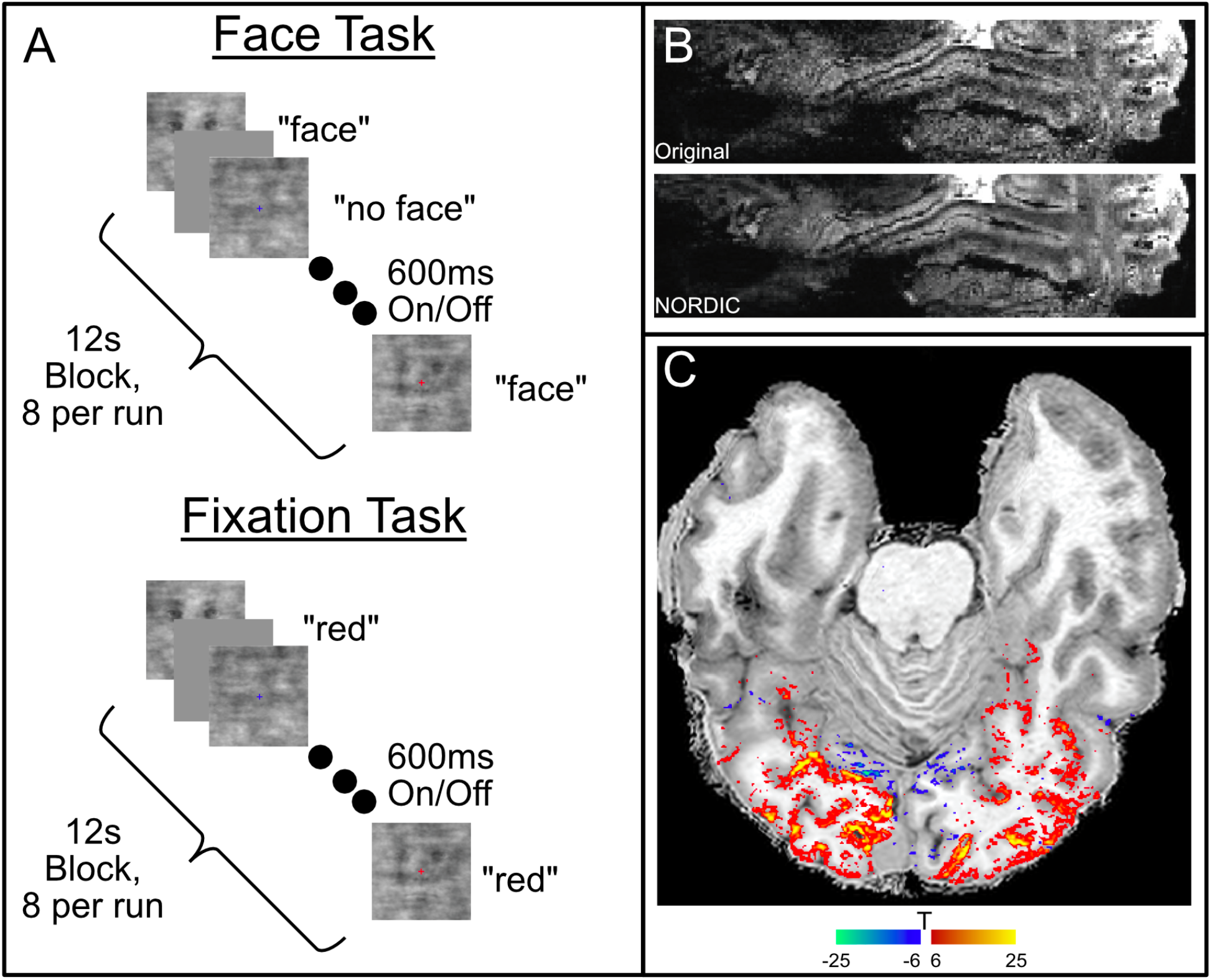
Paradigm, Coverage and Task Activation. **(A)** The schematic of the task design, consisting to 6 runs each of two tasks, which used identical stimuli and timing. Each run contained 8 pseudo-blocks, containing 10 stimuli (600ms ON/OFF) of faces of varying phase coherence. In the face task, subjects pressed a button to indicate whether they perceived a face or no face. In the fixation task, the subjects pressed a button when they observed the fixation point turn red. **(B)** example of single gradient echo EPI image from the fMRI time series before and after NORDIC denoising (sagittal orientation) illustrating coverage achieved for this study. **(C)** The mean activation for an exemplar subject (shown on the distorted coregistered T1 anatomical image) to the Face task shows robust activation across the brain, including areas distant from primary visual cortex.

Data were analyzed using a GLM to produce an estimate of activation in response to each individual block. The cortical ribbon was segmented in 3 equidistant depths using LayNii^19^ (Figure 2A) in the retinotopic representation of the stimuli in V1 (localized by combining probabilistic atlases^20^ and the activity elicited by the first block within each run, which was excluded from subsequent analyses), Occipital Face Area (OFA) and Fusiform Face Area (FFA) (identified by means of a face localizer – see Methods). Following NORDIC denoising, we found robust activation to faces throughout the image volume (Figure 1C), despite the intrinsically low native SNR of the high-resolution images. Furthermore, we found consistently larger activation in response to the face relative to the fixation task across all layers and ROIs (Figure 2C). These effects were largest in the FFA. Unsurprisingly, all areas also showed the expected pial bias inherent to gradient echo BOLD imaging^21–23^ (Figure 2C, 3 and 4).

**Figure 2.**
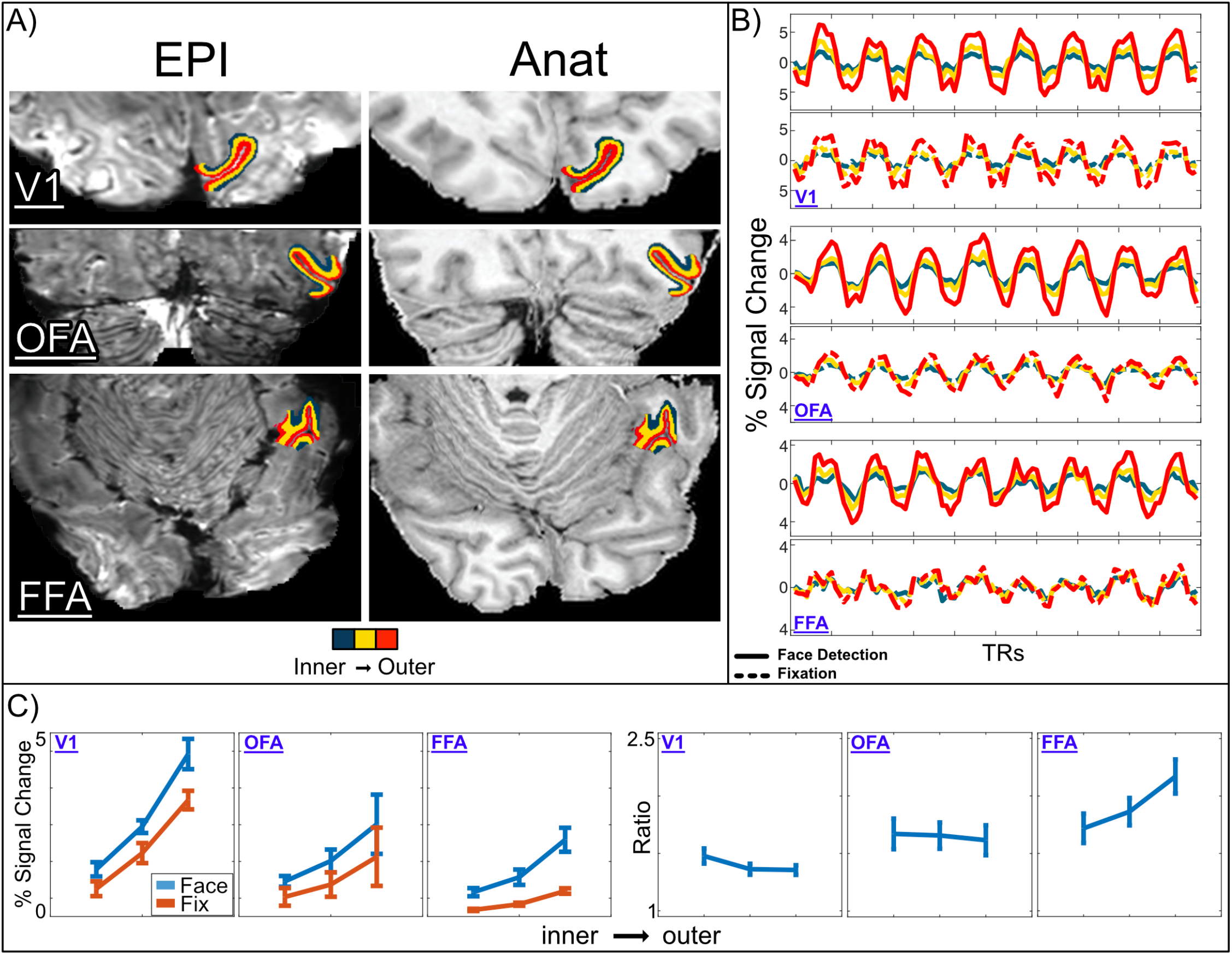
**(A)** The native space (i.e. still distorted due to B0 inhomogeneity) echo planar image (EPI) and anatomical image with distortion applied are shown. The 3-layer ROIs for V1, OFA and FFA are shown overlaying both. **(B)** Corresponding Single run timecourses for each task, depth and region. Cortical depths are depicted in different colors: Green for inner; Yellow for middle; and Red for outer depths **(C)** left, average percent signal change responses for all tasks, depths and regions. Error bars show standard errors across subjects. Right, average bootstrapped population across participants for the ratio between tasks (error bars show the mean of the 68% bootstrapped CIs). This figure is for descriptive purpose only as inferential statistics were carried out within participants (see Figure 4).

**Figure 3.**
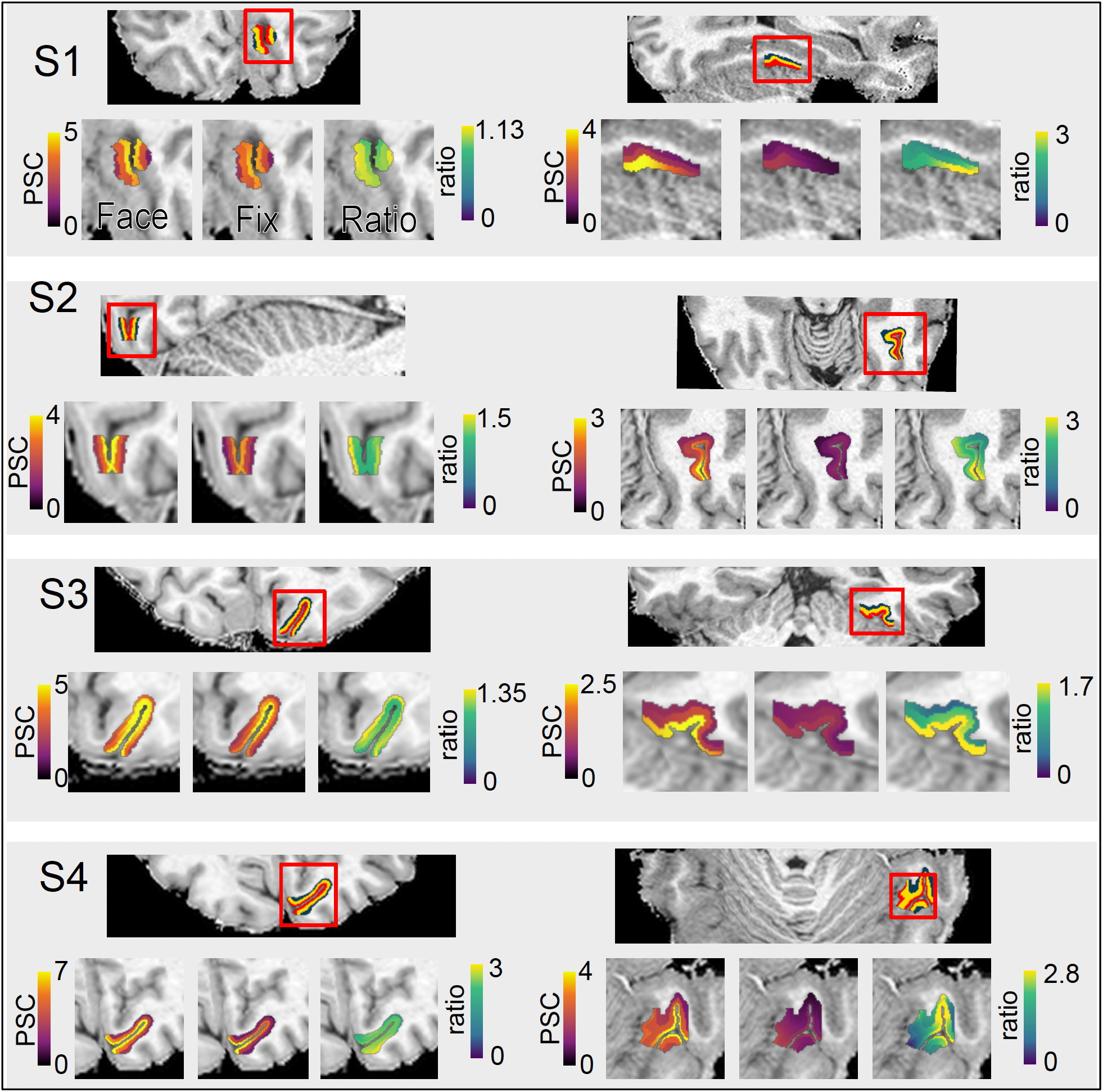
Layer-wise plots of Face, Fixation, and the Ratio of activation. For each subject we show the V1 (Left side) and FFA (Right side) ROI, with the 3 layer profiles shown (red being the outer, yellow the middle and green the inner depths). The zoomed-in portions show the precent signal change BOLD activation for face (left) and fixation(middle) task and their ratio (right). For visualization only, we performed within-layer smoothing (FWHM 3mm). In all subjects we observe the expected pial bias. In addition, we observe that the pattern of the ratio between tasks tends to increase towards inner layers within V1, and in contrast, the ratio between tasks increases towards outer layers in the FFA.

**Figure 4:**
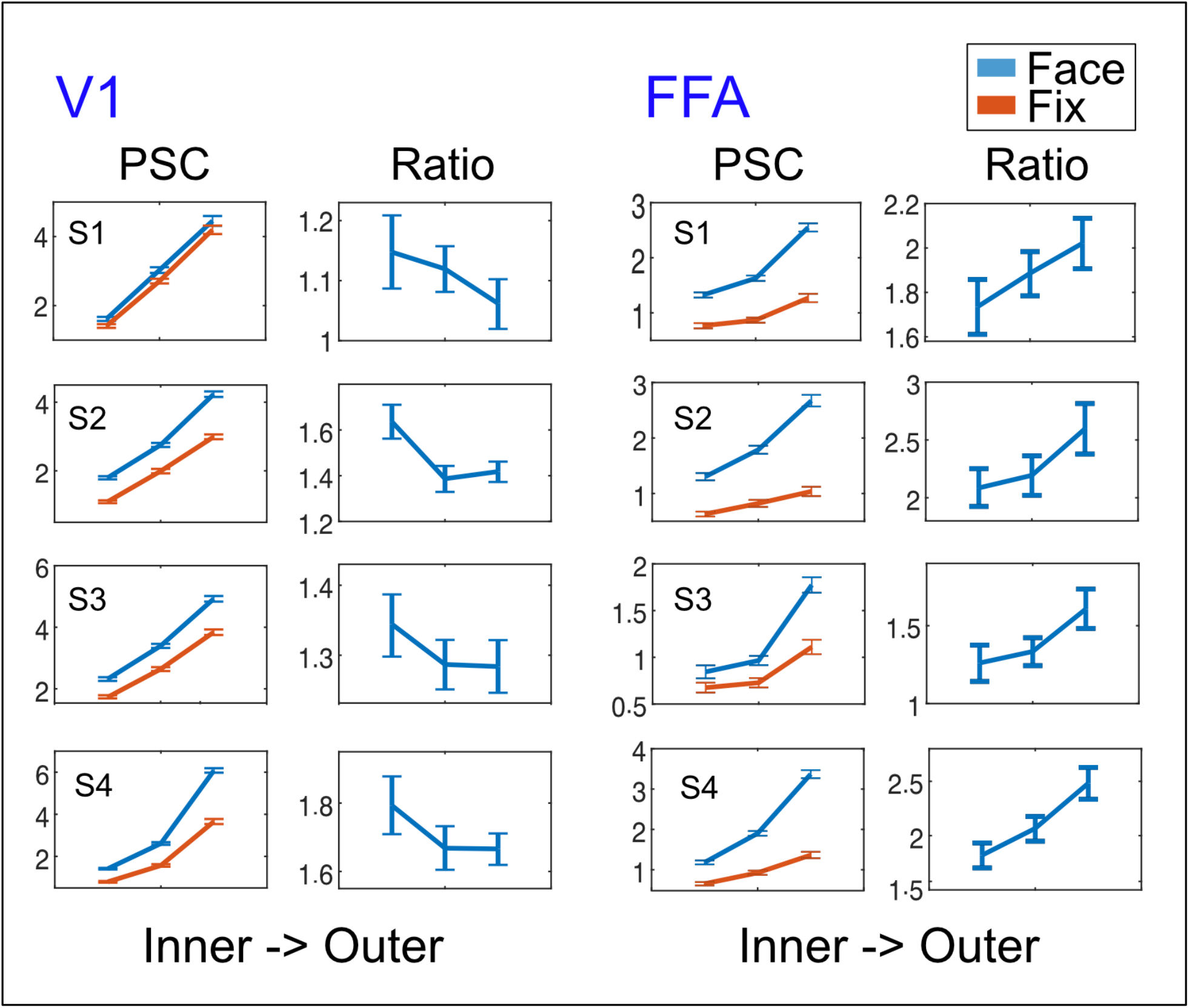
Percent Signal Change (PSC) and Ratios. Left two Columns: The 1st column shows the PSC for the Face (blue) vs Fixation (Orange) task within V1, with the expected pial bias. The 2nd column shows the ratio between the task, with the inner layers having a significantly higher ratio relative to the outer layers. Right two columns: Organization is as before, but here we observe the inverse pattern in the ratio, as the outer layer of the FFA is significantly larger than the inner layers.

To quantify task induced top-down modulations and account for the pial bias, we examined the layer-wise ratio of the activity elicited by the face detection over that elicited by the fixation task within our 3 ROIs. Ratio is not directly impacted by the BOLD magnitude of single conditions, which varies with vasculature; as such it is a more useful and interpretable parameter^24^. 95% bootstrapped confidence interval on the difference of the ratio across inner and outer depths showed that the ratio between tasks was largest (p<.05 Bonferroni corrected) in magnitude at the inner layers of V1 (95% Bonferroni corrected bootstrapped CI: S1 [0.03, 0.11]; S2 [0.09, 0.24]; S3 [0.01, 0.06]; S4 [0.05, 0.30];) and in the outer layers of the FFA (95% Bonferroni corrected bootstrapped CI: S1 [−0.39, −0.92]; S2 [−0.49, −1.09]; S3 [−0.38, −1.05]; S4 [−0.87, −1.31] - Figure 2 and 3). No statistically significant differences were observed in the OFA.

In this work we demonstrate, reliably within individuals, the feasibility of simultaneously imaging cortical depths across several areas along the visual hierarchy during complex human behavior. Cortical depth dependent fMRI is rapidly gaining momentum; however, the interpretation of findings can be challenging. Traditionally, comparisons against animal models have provided some assurance with regards to the validity of cortical depth dependent results. However, these validations are not always feasible nor desirable, as meaningful deviation between animal and human brain functions are expected, especially when examining higher order cognitive processes. With the approach employed here, we show that changes in the relative contribution of top-down signals (modulated via varying task demands) to different cortical depths can be used to assess hierarchical relationships across cortical areas, while addressing some of the main challenges of laminar fMRI. We further provide neuroscientific insights into the microcircuitry of conscious perception. These points are discussed in detail below.

Continued technological developments subsequent to the launch of 7Tesla magnets^25^ have led to the reliable acquisition of functional images with submillimeter precision, a feat primarily enabled by the ultra-high field strength. Undeniably, one of the main advantages of these ultra-high-resolution images is the ability to study mesoscale functional organization in humans non-invasively. At present, the precisions achievable with most pre-clinical invasive techniques remain unattainable with non-invasive human fMRI. However, such invasive techniques are not routinely applicable for human studies and, given the complexity of the human brain, it cannot be *a priori* assumed that animal studies provide accurate inferences on human brain function. In addition, fMRI affords larger cortical coverage, offering the unique possibility of studying neural processing and circuitry across distal areas differing in hierarchical complexity. However, like all techniques employed in the study of the brain, fMRI also has challenges.

Two of the major challenges faced in ultrahigh resolution human fMRI studies are the low SNR – especially in regions of the brain further away from the coil elements of the receiver array – and the difficulty of imaging at ultrahigh resolutions over large fields-of-views (FOVs). Consequently, ultrahigh resolution studies typically employ a highly restricted cortical coverage^8,26–28^ and have thus far focused on a single or closely located regions, limiting the impact and interpretations of their findings. In this work, we target multiple cortical regions in different, relatively distal locations across the visual hierarchy, some of which (such as the FFA) are associated with notoriously low SNR. To obtains this larger FOV and achieve adequate coverage requires accelerated acquisitions, especially if high spatial resolution is desired, leading to a further decrease in SNR due to k-space undersampling. We overcame these SNR challenges with the use of NORDIC^16,17^ (Figure 1B) denoising, which suppresses thermal noise (see Figure 2B), the dominant noise source in such ultrahigh resolution images. The resulting SNR was sufficient to permit*, within each participant*, the reliable identification of the stimulus locked activity elicited by each single block, across cortical depths, regions and tasks. This allowed performing statistical analyses across the 42 single blocks/trials within single subjects (with subjects representing 4 test re-test cases).

Another challenge in fMRI studies arises from the fact that the functional maps are a proxy for, rather than a direct measure of, neuronal activity. They are mediated by neurovascular coupling and the mechanisms involved in converting hemodynamic alterations to MR detectable signals. A prominent example where this complexity introduces obfuscations is the pial bias – i.e. the presence of apparent, albeit non-neuronal and therefore undesirable, functional responses originating from large draining vessels located on the pial surface. This problem is particularly pronounced in gradient echo (GE) EPI based functional images^23,29^, which is by far the most commonly used fMRI approach. While there are other acquisition techniques^29–32^ that are less sensitive to large vessels, we chose the commonly employed GE-BOLD because it affords higher functional contrast-to-noise ratio (fCNR) and SNR. Given the goals of this work, namely imaging cortical depths at even higher resolutions concomitantly across distal regions, capitalizing on the highest achievable fCNR and SNR becomes even more crucial. However, the GE related pial bias means that, despite the observed laminar correlates of top-down and bottom-up connections, interpretation of depth dependent fMRI data, as it pertains to neuronal processing within laminae, remains challenging, as differences in magnitude across depths are confounded by cortical vascular architecture. We overcame this limitation by comparing the activation elicited by multiple tasks within layers, across multiple brain regions simultaneously.

We assessed task-related modulations to identical visual stimuli across three areas with known differences in hierarchical position: V1, a low-level area receiving thalamic input and providing input to a variety of visual cortical areas; the OFA, a mid-level area located in inferior occipital gyrus associated with the processing of faces; and the FFA, a higher-level visual area also associated with face processing. We observed that the relative magnitude of task-related modulations across the cortical depths depends on hierarchical location: they were most pronounced in the inner depths of V1; not present in the OFA; and most pronounced in the outer depths of the FFA (Figures 2, 3 and 4). These differences are ascribed to the top-down contributions based on the tasks that generated them, as discussed further on.

The use of multiple tasks allows the examination of task-induced differences on BOLD responses *within* cortical depths and then compares the magnitude of these differences (rather than the magnitude of the responses themselves) across depths. In other words, in addition to assessing top-down modulations, contrasting the activation elicited by one task *versus* another also serves to normalize the pial bias related increase in fMRI signal magnitude across depths. Moreover, the observed differences in the laminar patterns of top-down modulations in different areas (e.g. inner dominant in V1 *versus* outer dominant in FFA) represents further assurance that our results are not a simple byproduct of the undesirable pial bias.

Comparing the activation across tasks to identical stimuli is crucial for isolating top-down modulations. Pure thalamic feedforward projections of identical visual input should lead to identical cortical responses. Therefore, meaningful differences in functional responses to identical stimuli across tasks can only be related to some top-down signal. However, interpreting amplitude differences across cortical depths while examining responses to a single task within a single region is challenging, especially if the observed effects are most pronounced in outer layers. A recent study^33^ employing a single task reported amplitude differences between expressive faces to be most pronounced in the outer depths of V1. While the authors ascribed their findings to top-down processes, they acknowledge that it is especially difficult to disentangle feedforward from feedback related signals with just a single task. Unlike the images used in the current studies – identical across tasks – the visual stimuli in ^33^ in fact greatly differed across the compared conditions. Given these experimental settings, it is thus impossible to exclude that their finding is not related to differential amounts of feed-forward thalamic activation rooted in low-level visual differences in the stimuli.

It would be plausible though, to expect that different types of feedback processes may target different layers. For example, the “higher level feedback” from distal regions (such as the one studied here) may terminate in the inner layers of V1, as opposed to “lower level feedback” from short range connections, which may be terminating in superficial layers in V1 instead^34^. Accordingly, a number of studies have reported inner (e.g. ^34–36^) or outer^37^ layer top-down modulations. However, without appropriate controls – such as varying task demands to identical visual inputs or the occlusion paradigm^38^ – the interpretation of these results remains difficult.

Our finding of larger task-related modulations in the outer layers in FFA (Figure 2, 3) is consistent with two theories: that of apical dendritic amplification^39^ in which an already firing neuronal population receives further amplification from higher level areas to apical dendrites; and with the notion of the FFA as an intermediate stage of processing^14,40^. This effect of amplified dendritic activity has been associated with conscious perception, increasing the flexibility of neural coding by rendering it sensitive to context^41^. It has further been suggested that this mechanism of apical amplification contributes to holistic integration of information^41^, a crucial process in face perception^42^. Accordingly, the FFA may be receiving feedback, in the outer layers, from a higher level face processing region, such as the anterior inferior temporal cortex, which appears to be highly sensitive to the perceived faceness of stimuli^14^.

Marvan et al.^41^ argue that perceptual content represented in the activity of pyramidal cells located in deep layers remains “unconscious” until it is amplified by apical amplification and that this mechanism can explain differences in the magnitude of activation to identical stimuli based on their perceptual status, reported here (Figures 2,3 and 4) and elsewhere^14,43^. While to directly test this hypothesis we would require, at a minimum, faster sampling to be able to infer the temporal evolution of top-down effects across layers, it is in line with what is reported in this work. The larger top-down task-related modulation in the outer depths of the FFA may be reflecting apical amplification and may therefore be responsible for the increased activity observed for the face relative to the fixation task.

The results of larger top-down modulations in the outer depths of FFA and in inner depths of V1 is also consistent with the reverse hierarchy theory^44,45^. The outer layer, context specific top-down modulations in the FFA (resulting from apical amplification), would then be fed back to inner depths of V1, where pyramidal cells are located and somatic integration occurs^41^. This would serve to “select” the relevant low-level information stored in early sensory areas (in this case V1) that will subsequently be fed-forward to higher level areas to fulfill task demands. Moreover, because the relative contribution of top-down effects is highly dependent on cortical depth, our results suggest that analyses that average over laminae and/or large volumes are likely to considerably underestimate the magnitude of top-down modulation on specific neuronal subpopulations.

Moving forward, the feasibility of further increases in the spatial resolution of fMRI will continue to facilitate human mesoscale studies. If we consider that the thickness of the cortical ribbon in V1 (where the cortex is thinnest) here measures 2.59 mm on average over subjects (with the minimum being 2.13 mm), and the highly folded nature of the cortex, it becomes evident how resolutions of 0.8 mm^3^, often used for laminar fMRI studies, may not suffice to fully characterize subtle response modulations across laminae. While our usage of 0.7mm^3^ voxels is only a 0.1mm reduction along each voxel dimension, it does represent a ~33% reduction in voxel volume (0.343 μL vs 0.512 μL) and therefore better samples the cortical depth. However, further increases in resolutions will be invaluable, as was reflected in the US Brain Initiative’s challenge to the community to move towards higher resolutions to better characterize fine neural microstructures^46,47^.

## Conclusion

In this work we showcase the potential of laminar-fMRI to explore large scale network activity along distant regions of differing complexity along a processing hierarchy, while further pushing the spatial resolution of functional images. We find that even a relatively simple behavioral manipulation is able to uncover distinct activity profiles, which is to be expected from the vastly different computations occurring in V1 and the FFA. While cortical-depth-dependent (or layer-) fMRI is rapidly gaining popularity because of its immense potential, the interpretation of depth-dependent findings can be challenging due to a number of factors – including, but not limited to, the complexities of neurovascular coupling, the high, albeit still limited spatial resolution and the low SNR. The concerns are especially salient when deviation from animal results are observed. Validation against animal models is not always possible nor desirable, as meaningful differences with the human brain are expected, especially when examining higher order cognitive processes. We argue that the approach employed here, entailing the use of several tasks and imaging different distally located areas, constitutes a promising path towards characterizing laminar microcircuitry in humans during complex behavior. Our approach further allows drawing comparisons with electrophysiological and even neurobiological results and may provide support for broader neuroscience theories. As methodologies continue to evolve, it is conceivable that whole brain, layer-wise activation profiles could be explored at even higher resolutions, in order to fully realize the potential of cortical depth dependent fMRI and better understand the mesoscopic organization of the human brain.

## Methods

### Participants

A total of 4 participants were scanned for this project. All subjects had normal, or corrected vision and provided written informed consent. The local IRB at the University of Minnesota approved the experiments.

### MRI Scanning

All functional MRI data were collected in a single session with a 7T Siemens Magnetom System using a 1 by 32-channel NOVA head coil. T2*-weighted images were collected using sequence parameters (0.7mm isotropic voxels, TR 2.03s, Multiband 2, GRAPPA 3, 6/8ths Partial Fourier, TE 29.6ms, Flip Angle 65°, Bandwidth 1050Hz). For each participant we manually shimmed the B0 field to maximize homogeneity over lateral occipital and ventral temporal regions.

In addition, we also performed manual shimming of the B0 magnetic field prior to data collection, in order to reduce inhomogeneity within the right ventral temporal cortex. Additional processing steps risk the potential to produce detrimental spatial blurring.

T1-weighted anatomical images were obtained using an MP2RAGE sequence (192 slices; TR, 4.3s; FOV, 325 x 240mm; flip angle, 4°; TE 2.27ms; spatial resolution, 0.75mm isotropic voxels) which were collected halfway through the same scanning session. Timing was chosen to reduce participant fatigue. Background noise in the unified MP2RAGE image was removed^48^ using code implemented in LayNii (LN_MP2RAGE_DNOISE). Anatomical images were used only to supplement region of interest (ROI, see Regions of Interest below) drawing and for visualization purposes.

### Primary Tasks and Stimuli

We used grayscale images of faces (20 male, 20 female). We manipulated the phase coherence of each face from 0% to 40% in steps of 10%, creating 160 images. Stimuli subtended approximated 9 degrees of visual angle. Each image was cropped to remove external features using an eclipse. Eclipses spanned the full vertical extent of each face and 80% of the external extent. The edge of the eclipse was smoothed by convolution with an average filter (using “fspecial” with “average” option in MatLab). This procedure was implemented in order to prevent participants from performing simple edge detection using the hard edges typically present in face images.

The amplitude spectrum of each image was equated and we controlled the Fourier spectra across stimuli, ensuring that the rotational average amplitudes for a given spatial frequency were equated across images while preserving the amplitude distribution across orientations ^49^. The standard deviation of pixel intensity was also kept constant across stimuli.

These images were used in a pseudo-block design for the 2 tasks, with 10 stimuli presented in a 600ms on/600ms off for a total block duration of 12 seconds with an inter-block interval of 12 seconds. For each task run 8 blocks were shown, with an additional 12s “off” period at the beginning of the scan producing a total run length of 204s. For each task, the 6 runs were collected, for a total of 12 runs associated with the primary tasks. During all tasks, the fixation cross remained black during the off period, but cycled (at 4Hz) between red, green, blue and yellow during the stimulus on periods.

For each run of the primary tasks participants were instructed to maintain fixation on a centered crosshair throughout the entire run and to perform one of 2 tasks, similar to our prior work^14^. These were 1) a domain-specific, stimulus-relevant task, requiring the perceptual judgment of the visual stimulus and 2) a non-specific, stimulus-irrelevant attention task. The domain-specific “Face Detection” task required that subjects maintain fixation and respond to each individual stimulus indicating whether they perceived a face or did not perceive a face. Instructions were carefully delivered to inform the subjects that there were no correct answers, as we were interested only in their subjective perception. For the non-specific attention “Fixation Detection” task, which used identical stimuli, participants were instead required to report when the fixation cross turned to a red color. As this occurred only during the ‘stimulus on’ periods, the pattern of responses was matched between the 2 tasks, despite different overall task demands. Visual stimuli were identical, their sequential presentation within a block and cross hair timing was matched across tasks to ensure that any differences observed were related solely to topdown processes. Tasks were blocked by run and counterbalanced across participants.

### Localizer Task

During the same scanning session we also completed 3 runs of a face localizer task. Each localizer run consisted of blocks of faces, objects and noise textures. Each run began with a 12s “off” period, with a black fixation cross displayed on a grey background for 12 sec. Then, 9 randomly presented blocks of images were shown, with each block (3 blocks/category; separated by a 12 sec fixation) involving the presentation of 10 different stimuli randomly presented for 800 ms, separated by a 400 ms ISI. We implemented a 1-back task, in which subjects were instructed to respond by pressing a button when a stimulus was repeated, which occurred on about 10% of the trials. Each block occurred 3 times within a run, for a total run duration of 228 seconds.

### Data Processing

The DICOM files were converted to separate magnitude and phase images using dcm2niix^50^. Prior to any other processing, we applied the NORDIC denoising technique to suppress thermal noise in the repeated image timeseries^16,17^. Next, we applied conventional processing steps using AFNI version 20.3.05^51^, which included slice timing correction, motion correction, and alignment to a functional reference image. The functional reference image was chosen as the first image from the run prior to the anatomical image acquisition. Alignment to the functional reference image was initiated using AFNI tools, with an affine transform between the mean of each run timeseries and the mean of the motion corrected run containing the reference image. An additional nonlinear transform was calculated (ANTs version 2.3.5) using the BSplineSyN^52^ warp with cross correlation metric between each run and the mean of the reference run. All transforms (motion, affine, nonlinear) were combined and carried out in one step to minimize interpolation-induced blurring and preserve spatial specificity. Upsampling was performed simultaneously in this step, writing out each final and fully aligned image with to a 0.35mm grid to preserve fine scale spatial information that would typically be lost due to interpolation^18^.

The corresponding single band reference image for reference image run, as well as the nearest reverse phase single band reference image were used to calculate a warp field required for distortion correction. These single band reference images were used due to their higher contrast^53^. A temporary warped reference image was then aligned to the anatomy using an affine transform (3dAllineate). As we chose to stay in distorted image space to reduce blurring or information loss, we then inverted the epi-to-anatomical transform, as well as the distorted- to-undistorted space warp field. These transforms were then combined into a single step and applied to the anatomical image with simultaneous upsampling, producing a 0.35mm anatomical reference image in the original functional image space.

### Regions of Interest

Regions of interest were manually drawn using itk-SNAP^54^. In total, 6 ROIs were drawn, left and right primary visual cortex (V1), occipital face area (OFA) and fusiform face area (FFA). The ROIs were drawn directly on the functional image in distorted space, using the mean functional image (across all runs) as the primary reference. In addition, the distorted anatomical image was also used as an additional guide for grey/white/CSF boundaries. In all cases, the functional image served as the definitive reference, particularly in areas affected by signal dropout, and layers were restricted to gray matter, creating ROIs that were faithful to the functional data.

The V1 ROI was based on combining information from probabilistic atlases^20^ and the mean of the first block from each run (including both tasks), which was subsequently discarded from further analyses. The OFA and FFA ROIs were drawn based on the Faces>Noise & Objects contrast (see below). ROIs were defined with 3 values, marking gray matter, white matter boundary and CSF boundary. These were converted to 3 equidistant layer profiles using LayNii. These layer ROIs were then used in subsequent analyses.

The increased resolution also requires further attention to layering analysis methods, as automated solutions may not be effective. Here we drew layers manually using the EPI images as a reference. As the EPI contrast can be challenging to use as a layer guide, we also used T1 weighted anatomical images, however we reversed the direction of typical processing. In the current work we used displacements maps (computed using images acquired with reverse phase encoding polarity – see data preprocessing) to distort the T1 to match the functional image space. This avoids large scale nonlinear image warping, and thereby helps preserve spatial specificity^18,55^. Moreover the right left phase encode direction was chosen to minimize distortions in FFA and V1, the primary regions of interest in this study. Distorting the T1, rather then correcting the EPI images further reduces the number of interpolation steps, therefore preventing unnecessary unwanted spatial blurring.

### Data Analysis and Modelling

The upsampled data corresponding to the 2 primary tasks as well as the localizer was loaded into MatLab for further data analyses. For the 2 primary tasks, we constructed a general linear model (GLM) to calculate parameter estimates (betas) and corresponding t-statistics. This was done by modeling each pseudo-block as single event, with duration 12s, to create estimates for each individual “block”. These betas were then converted to percent signal change (PSC) values and used in further analyses. For the functional localizer, we constructed a GLM to evaluate activation across all runs to the blocks of Faces, Objects and Noise categories. Betas and t-statistics for the face contrast, as well as for Faces > Average(Noise + Objects) were used to guide ROI drawing. For additional guidance during the ROI drawing process (see above), we also repeated the localizer model with 1 (0.7mm) and 2 (1.4mm) voxel FHWM Gaussian smoothing. Note, that no smoothing was applied to the primary task data used for analysis; as smoothing was used only to aid in identification of ROIs.

### Statistical Analyses

The PSC values were then extracted for each ROI on a layer-by-layer basis. We performed the 20% trimmed mean, averaging across voxels within each layer and performed the ratio across tasks between the responses elicited by the matched blocks of stimuli. To infer statistical significance, we performed Bonferroni corrected 95% bootstrapped confidence intervals on the ratio across tasks for the difference between innermost and outermost depths. 2 ways bootstrapped confidence intervals were computed as follows. For a given ROI, we performed the trial-by-trial ratio difference between innermost and outermost depths. We therefore sampled with replacement the 42 ratio differences, performed the mean, and stored that value. The procedure was repeated 10000 times. We sorted these values in ascending order and selected the ~99.166 and ~0.083 percentiles (respectively corresponding to the .975 and .25 percentiles Bonferroni corrected by the number of tests, in this case 3). This procedure was repeated for each of our 3 ROIs. Confidence intervals not including zeros were considered to be statistically significant at p<.05 (Bonferroni corrected).

## ACKNOWLEDGEMENTS

This work was supported by NIH grants P41 EB 027061 (Ugurbil), U01 EB025144 (Ugurbil) and RF1 MH117015 (Ghose), RF1 MH116978 (Yacoub)

